# *Artemisia* extracts Differ from Artemisinin Effects on Human Hepatic CYP450s 2B6 and 3A4 *in vitro*

**DOI:** 10.1101/2022.06.24.497548

**Authors:** Ndeye F. Kane, Bushra H. Kiani, Matthew R. Desrosiers, Melissa J. Towler, Pamela J. Weathers

## Abstract

**Ethnopharmacological relevance:** The Chinese medicinal herb, *Artemisia annua* L., has been used for >2,000 yr as traditional tea infusions to treat a variety of infectious diseases including malaria, and its use is spreading globally (along with *A. afra Jacq. ex* Willd.) mainly through grassroots efforts.

**Aim of the study:** Artemisinin is more bioavailable delivered from the plant, *Artemisia annua* L. than the pure drug, but little is known about how delivery via a hot water infusion (tea) alters induction of hepatic CYP2B6 and CYP3A4 that metabolize artemisinin.

**Materials and Methods:** HepaRG cells were treated with 10 μM artemisinin or rifampicin (positive control), and teas (10 g/L) of *A. annua* SAM, and *A. afra* SEN and MAL with 1.6, 0.05 and 0 mg/gDW artemisinin in the leaves, respectively; qPCR, and Western blots, were used to measure CYP2B6 and CYP3A4 responses. Enzymatic activity of these P450s was measured using liver microsomes and P450-Glo assays.

**Results:** All teas inhibited activity of CYP2B6 and CYP3A4. Artemisinin and the high artemisinin-containing tea infusion (SAM) induced CYP2B6 and CYP3A4 transcription, but artemisinin-deficient teas, MAL and SEN, did not. Artemisinin increased CYP2B6 and CYP3A4 protein levels, but none of the three teas did, indicating a post-transcription inhibition by all three teas.

**Conclusions:** This study showed that *Artemisia* teas inhibit activity and artemisinin autoinduction of CYP2B6 and CYP3A4 post transcription, a response likely the effect of other phytochemicals in these teas. Results are important for understanding *Artemisia* tea posology.

## 1. Introduction

Artemisinin (ART), produced by the Chinese medicinal herb, *Artemisia annua* L. (Hsu, 2006), is extracted and chemically derivatized to more bioavailable forms including artesunate, artemether, and dihydroartemisinin for incorporation into the current antimalarial artemisinin combination therapies (ACTs) (WHO, 2022). Although ACTs are widely distributed, their costs and accessibility are problematic (O’Connell et al., 2011), so many people still use the >2,000 yr traditional *Artemisia sp*. tea infusions to treat a variety of infectious diseases including malaria (Hsu, 2006). Furthermore, *A. annua* use is spreading globally (along with *A. afra* Jacq. *ex* Willd.) mainly through grassroots efforts, and thus it is important to compare traditional and modern pharmacology of ART to improve our understanding of ART delivered via the plant (Feng et al., 2020).

Cytochrome P450s (CYPs) are essential in first-pass drug metabolism. These enzymes can be induced or inhibited by drugs and can interfere also in the therapeutic response of many drugs (Lynch and Price, 2007). There are many hepatic P450 enzymes, and two in particular, CYP3A4 and CYP2B6, are known to metabolize artemisinin (ART; Fig. 1) (Lee and Hufford, 1990; Svensson and Ashton, 1999). In the liver, ART is metabolized mainly through CYP2B6 with a minor contribution from CYP3A4 to yield deoxyartemisinin, crystal 7, deoxydihydroartemisinin, and 9,10-dihydrodeoxyartemisinin (Lee and Hufford, 1990; Svensson and Ashton, 1999), all of which are therapeutically inactive against malaria. CYP3A4 and CYP2B6 are active in human liver microsomes (HLMs), primary human hepatocytes (PHHs), and HepaRG cells. Both PHHs and HepaRG cells are used for transcript induction and inhibition studies of P450s, with HepaRG better suited for routine testing (Bernasconi et al., 2019). PHHs have long been limited by availability, and are subject to variability among individual donors (Bernasconi et al., 2019; Costa et al., 2014). HepaRG cells provide an immortalized hepatocyte human cell line that provides better testing consistency (Aninat et al., 2006; Bernasconi et al., 2019; Le Vee et al., 2006).

**Figure 1.**
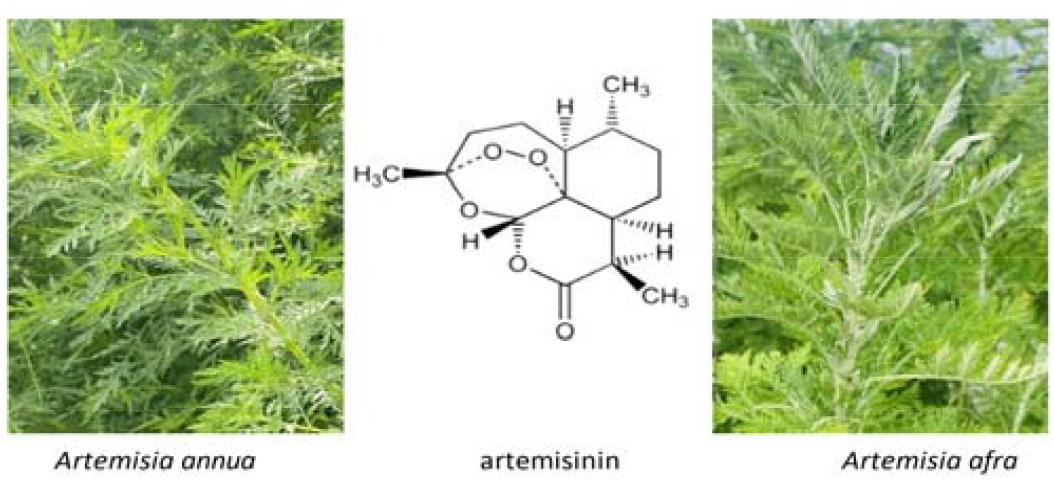
Artemisinin, *Artemisia annua*, and *A. afra*.

Tea infusions have been the traditional means for >2,000 yrs whereby *A. annua* was consumed to treat fever-generating ailments, but there have been concerns raised by the World Health Organization (WHO, 2019) that continued consumption of these daily prepared teas would deliver diminishing amounts of ART because of ART’s P450 metabolism in the liver and its known auto-induction of CYP 2B6 and CYP3A4 in PHHs (Elsherbiny et al., 2008; Simonsson et al., 2003; Xing et al., 2012) and *in vivo* (Asimus et al., 2007). When ART is delivered as the traditional hot water extract (tea infusion) of dried leaves of *A. annua* (DLA), the bioavailability of ART increases >40-fold (Desrosiers et al., 2020; Weathers et al., 2011; Weathers et al., 2014), suggesting that other compounds present in DLA intervene to inhibit ART metabolism in the liver. Indeed, along with improved ART solubility from DLA’s essential oils (Desrosiers et al., 2019; Desrosiers and Weathers, 2016) and enhanced intestinal transport shown with Caco-2 cells (Desrosiers and Weathers, 2018), inhibition of hepatic CYP3A4 and CYP2B6 together enhanced ART bioavailability (Desrosiers et al., 2020). While some phytochemicals found in *A. annua*, e.g., arteannuin B (AB) and quercetin, both inhibited CYP2B6 and CYP3A4, inhibition was not as potent as the extract *in toto*. Others also reported that AB and many other phytochemicals inhibited CYP3A4 (Ashour et al., 2017; Cai et al., 2017; Desrosiers et al., 2020; Wei et al., 2015; Zehetner et al., 2019) and even CYP2B6 (Desrosiers et al., 2020). The Desrosiers et al. (Desrosiers et al., 2020) study showed that another related species, *A. afra*, used indigenously in Southern Africa to treat malaria and other infectious diseases (Du Toit and Van der Kooy, 2019), also inhibited both P450s. Compared to the results from *A. annua* with ∼1.4% ART content, *A. afra* had <0.05% ART, so results suggested that it is important to also measure the effect of *A. afra* extracts on hepatic induction of these P450s. To our knowledge there are no studies measuring the effect of *Artemisia sp*. tea infusions on induction of either CYP2B6 or CYP3A4.

The inhibition of CYP3A4 and CYP2B6 is usually undesirable because it could lead to drug toxicity, drug-drug and herb-drug interactions, and other serious adverse effects. However, when carefully managed, inhibition also can improve the therapeutic efficacy of drugs (Samuels and Sevrioukova, 2021). A recent clinical study showed *A. annua* extracts were safe and improved liver function (Han et al., 2020). In our study, hot water extracts of three different *Artemisia sp*. with ART contents ranging from 0-1.6% (w/w) were compared to ART for their effect on transcription, translation, and enzymatic activity of CYP2B6 and CYP3A4. Our results improve our understanding of the role of those medicinal plants, delivered as DLA, on first pass liver metabolism and guide future investigations into herb-drug interactions that are crucial for safe use of medicinal herbs (Wanwimolruk et al., 2014; Wanwimolruk and Prachayasittikul, 2014).

## 2 Materials and Methods

### 2.1 Biologics, Chemicals, and Reagents

Dried leaves of *Artemisia annua* L. cv. SAM (voucher MASS 317314) were obtained from Atelier Temenos LLC in Homestead, FL (B#1.Sh6.01.15.20). Dried leaves of *Artemisia afra* Jacq. *ex* Willd. cv. SEN and MAL originated from Senegal and Malawi, respectively; SEN (voucher LG0019529) was gifted from Lucile Cornet-Vernet, and MAL was from Atelier Temenos LLC in Homestead, FL (voucher FTG 181107; B#1.RbA.10.12.20). HLMs were from Sekisui XenoTech (H2620); due to pandemic supply chain issues, HepaRG cells were purchased from Thermo Fisher (HPRGC10) or from Lonza (NSHPRG), along with the necessary media and supplements required for CYP induction, Western blots, and P450-Glo assays as follows: from Thermo Fisher: William’s E Medium (A1217601), GlutaMAX™ Supplement (32551020), HepaRG™ Thaw, Plate & General Purpose Medium Supplement (TPGP medium; HPRG670), HepaRG™ Serum-free Induction Medium Supplement (SFI medium; HPRG650), HBSS (14175-095), RIPA Lysis and Extraction Buffer (89900), DyNamo Flash SYBR Green qPCR Kit (F-415), Pierce™ Fast Western Blot Kit (35050), Pierce™ BCA Protein Assay Kit (23225), NuPAGE™ LDS Sample Buffer (4X) (NP0007), NuPAGE™ Sample Reducing Agent (10X) (NP0004), NuPAGE™ MES SDS Running Buffer (20X) (NP0002), Nitrocellulose Membranes (77010); from Qiagen: RNeasy Mini Kit (74106), RNase free DNase Set (79254), QuantiTect Reverse Transcription Kit (205313); from Promega: P450-Glo™ CYP3A4 assay (V9002), P450-Glo™ CYP 2B6 assay (V8322), NADPH regeneration system (V9510); from Sigma Aldrich: DMSO (D8418), rifampicin (R3501), quercetin (Q4951); from Cayman Chemical: artemisinin (11816), ketoconazole (15212); from Toronto Research Chemicals: clopidogrel (C576250); from AbCam: Recombinant Anti-Cytochrome P450 2B6/CYP2B6 antibody (ab140609); from ABclonal: CYP3A4 Rabbit pAb (A2544), HRP Goat Anti-Rabbit IgG (AS014), GAPDH (Human Specific) Rabbit mAb (AC036).

### 2.2 Artemisia leaf extraction and analyses

*A. annua* L. (SAM) and *A. afra* (MAL and SEN) hot water extracts were prepared as follows: 2 g dried leaves was added to 200 mL boiling water (equivalent to 10 g/L) on a stir plate and boiled for 10 min, then decanted through a 2 mm stainless steel sieve to retain most solids, cooled, and sterile-filtered (0.22 μm) before storage at -20°C. Artemisinin and total flavonoids were extracted from dried leaves by placing test tubes containing dried leaves and methylene chloride at a ratio of 25 mg to 4 mL in a sonicating water bath for 30 min, after which the solvent portions were dried under nitrogen flow and stored at -20°C until analysis. The level of artemisinin in the water extracts or dried leaves was determined using GC-MS according to the method detailed in (Martini et al., 2020). Authentic artemisinin was used as standard. Total flavonoid content was measured by the AlCl_3_ method of (Arvouet-Grand et al., 1994) with minor modifications (see Supplemental Material for details), and quercetin was used to generate a standard curve. ART and total flavonoid contents of the 3 cultivars of *A. annua* used in these experiments is shown in Table 1.

**Table 1.** Artemisinin (ART) and total flavonoid contents of *Artemisia* cultivars used in this study

|  |  |  |
| --- | --- | --- |
| *Artemisia* species & cv. | ART (mg/g DW) | Total flavonoids (mg/g DW) |
| *A. annua* SAM | 16.42 | 3.60 |
| *A. afra* SEN | 0.045 | 3.64 |
| *A. afra* MAL | nd | 6.84 |

DW=dry weight; nd=not detectable, but assigned a value of $1 \times 10^{-9}$ mg/g DW for later calculation purposes

### 2.3 CYP2B6 and 3A4 microsomal activity assays

P450-Glo assay kits for CYP2B6 and CYP3A4 were used to determine IC_50_ values using pooled (200 individuals) HLMs for all tea samples (SAM, MAL, and SEN) and pure ART according to the methodology of (Desrosiers et al., 2020) with some modifications. SAM tea samples were in the range of 0.28-145.42 μM ART, and MAL and SEN teas were in equivalent leaf dry weight amounts. DMSO was used as a solvent control at ≤0.25% (v/v).

### 2.4 Induction of CYP2B6, CYP3A4 enzymes in HepaRG cells

HepaRG cells were thawed at 37°C in a water bath, then 1 mL of suspended cells were aseptically transferred into a sterile 15 mL Eppendorf tube containing 9 mL of the pre-warmed HepaRG™ Thaw, Plate & General Purpose Medium (TPGP medium) resulting in a 1:10 ratio of cell suspension to total volume. Cells were washed and resuspended into the same medium for a total volume of 10 mL cells suspension then plated into a clear-bottom 24-well collagen-coated plate (for optimal cell attachment) according to the manufacturer’s protocol at a seeding density of ≅400,000 cells per well. After plating, the HepaRG™ cells were incubated at 37°C, 95% humidity with 5% CO_2_ for 6 h to allow recovery from freezing, then the TPGP medium was renewed, and cell morphology observed under a microscope. Cells were cultured for 72 h. At the beginning of the induction, TPGP medium was replaced by HepaRG™ Serum-free Induction Medium (SFI medium) to perform the 3-d induction test. SFI medium contained the following for the assay: 10 μM rifampicin (RIF) in 0.1% DMSO as positive control; *A. annua* (SAM) tea extracts with different concentrations of ART (10, 25, 50 μM); tea extracts of *A. afra* (SEN and MAL) at the equivalent biomass amount used to prepare *A. annua* SAM at 10 μM ART; and 10 μM pure ART in 0.1% DMSO. SFI medium containing 0.1% DMSO was used as the solvent control for RIF and ART, and medium containing distilled water was used as the solvent control for the tea extracts. Each SFI medium was replaced every 24 h for 3 d. HepaRG cultures were viewed and photographed daily to observe any changes in cell morphology after vehicle or drug addition (see Supplemental Material) (Xing et al., 2012). After the treatment period, cells were washed with HBSS twice before RNA extraction.

### 2.5 CYP2B6, CYP3A4 total RNA extraction and reverse transcript into cDNA

The total RNA from the hepatocytes was extracted according to the manufacturer’s instructions using the QIAGEN RNeasy mini kit. DNase I digestion was performed to avoid genomic DNA contamination and RNA determined by NanoDrop. Purity was indicated by the ratio of absorbance at 260 and 280. Total RNA (500 ng) was reverse transcribed using cDNA QuantiTect Reverse Transcription Kit then diluted 4-fold for polymerase chain reaction. A total reaction volume of 10 μL was made for each well in a 96 well PCR plate and analyzed to determine the CTs for each gene in the positive and negative samples, the mixture contained cDNA 25ng, 500 nM reverse and forward primers (for 3A4 and 2B6), 500 nM reverse and forward primer (for β-actin), and 1X DyNamo Flash SYBR green qPCR master mix. The program was 10 min at 95°C, 50 cycles of 15 s at 95°C, 1 min at 60°C, then 30 s at 72°C, followed by a dissociation curve step. β-actin was used as the reference gene, and the same primer sequence used by (Westerink and Schoonen, 2007) was used to amplify the targeted genes for 3A4 and 2B6: for 3A4 were (forward: CAG GAG GAA ATT GAT GCA GTT TT; reverse: GTC AAG ATA CTC CAT CTG TAG CAC AGT), for 2B6 (forward: TTA GGG AAG CGG ATT TGT CTT G; reverse: GGA GGA TGG TGG TGA AGA AGA G) and for β-actin (forward: CTG GCA CCC AGC ACA ATG; reverse: GCC GAT CCA CAC GGA GTA CT)

### 2.6 Gene expression

The expression of CYP2B6 and CYP3A4 after the assay was determined by qPCR. Primers were used to amplify the targeted genes using a Roche Light-Cycler 96 system. The mRNA levels of each test gene (2B6 and 3A4) were normalized to β-actin according to the following formula: CT (test gene) - CT (β-actin) = ΔCT. Thereafter, the relative mRNA levels of each gene were calculated using the ΔΔCT method: ΔCT (test gene in the treated sample) - ΔCT (test gene in the untreated sample) = ΔΔCT (test gene). The fold changes of mRNA levels were expressed as the relative expression 2^-ΔΔCT^. A fold induction greater than 2 relative to the vehicle (solvent) control and at least 20% relative to the positive control was defined as induction.

### 2.7 Western Blotting

HepaRG cells from Thermo Fisher at a seeding density of 400,000 cells/well were used for the Western blotting. Total proteins were extracted using RIPA buffer. Following extraction, protein was quantified using BCA assay following the manufacturer’s protocol (Thermo Fisher 23225). Western blotting was performed according to manufacturer’s protocol (Thermo Fisher 35050) with slight modifications. The proteins were separated on SDS-PAGE and transferred to nitrocellulose membrane. Protein electrophoresis was performed with 10% resolving and 4% stacking gels. Proteins were normalized, and 15 μg of protein with 1X NuPAGE LDS loading sample and 1 μL of NuPAGE sample reducing agent after heating at 70°C for 10 min was loaded into the gel. A Mini-PROTEAN® Tetra Cell (Bio-Rad) was used for protein electrophoresis at 100 volts to separate the proteins for 1.5 hr. After separation, proteins were transferred to the nitrocellulose membrane in transfer buffer (50 mL of NuPAGE transfer buffer 20X, 100 mL methanol, 1 mL NuPAGE antioxidant, and 850 mL distilled water) at 100 volts for 2 hr. Pierce™ Fast Western Blot Kit was used for membrane probing. Membranes were probed with anti-CYP2B6 and anti CYP3A4 primary antibodies. HRP goat anti-rabbit IgG was used as a secondary antibody. GAPDH (Human Specific) Rabbit mAb was used as the loading control. Immunoblots were developed by using chemiluminescence (Azure c600 gel imaging system).

### 2.8 Statistical Analyses

HLM P450 activity analyses had 2-3 technical replicates and IC_50_ values were calculated using GraphPad Prism 9.3.1 through nonlinear regression. Transcription analyses for each P450 had three replicates per batch of cells and SAM, ART and RIF were further replicated using 2-3 batches of cells. Induction was deemed significant at > 2-fold increase relative to the solvent control, and at least 20% relative to the positive control (Kenny et al., 2018). Western blots were at least twice replicated for each P450. Correlations between CYP450 activity and artemisinin or total flavonoids used Spearman’s Rho analysis (http://www.biostathandbook.com/spearman.html). Data were analyzed for significance using appropriate statistical tests.

## 3. Theory

ART in *Artemisia* extracts is more bioavailable and this study provides mechanistic evidence explaining hepatic P450 inhibition from transcription through translation and P450 enzyme activity in response to traditionally used *Artemisia* hot water extracts compared to ART.

## 4. Results

### 4.1 A. annua and A. afra tea extracts inhibit 2B6 and 3A4 enzyme activity

All three Artemisia sp. extracts inhibited the activity of both 2B6 and 3A4 with IC_50_ values ranging from 0.27-0.76 g/L of dried leaves for 2B6 and 0.31-0.34 g/L for 3A4 (Table 2). To calculate IC_50_s based on the ART content for all extracts, MAL, which had undetectable ART, was assigned an extremely low non-zero amount of ART (Table 1 footnote). ART also inhibited the activity of 2B6, but not 3A4. The correlation of IC_50_ to tea total flavonoid content using Spearman’s correlation (Fig. 2) was significant between 3A4 IC_50_ and total flavonoids (rho = 0.748 and p < 0.001) but was insignificant for 2B6 (rho = 0.157 and p = 0.533). Calculation of Spearman’s correlations for ART were not valid because there were no ART levels between the two extremes in these experiments.

**Table 2.** Inhibition of CYP2B6 and CYP3A4 activity by artemisinin and tea extracts in human liver microsomes.

|  | CYP2B6 IC<sub>50</sub><br>(μM ART) | CYP2B6 IC<sub>50</sub><br>(g DW/L)* | CYP2B6 IC<sub>50</sub><br>(mg FLV<sup>^</sup>/L) | CYP3A4 IC<sub>50</sub><br>(μM ART) | CYP3A4 IC<sub>50</sub><br>(g DW/L)* | CYP3A4 IC<sub>50</sub><br>(mg FLV<sup>^</sup>/L) |
| --- | --- | --- | --- | --- | --- | --- |
| **ART** | 7.51 | n/a | n/a | >600 | n/a | n/a |
| **SEN tea** | 0.09 | 0.58 | 87.87 | 0.05 | 0.34 | 51.72 |
| **MAL tea** | 2.69x10<sup>-9</sup> | 0.76 | 258.8 | 1.20x10<sup>-9</sup> | 0.34 | 115.5 |
| **SAM tea** | 15.46 | 0.27 | 104.8 | 18.13 | 0.31 | 122.9 |

\* = dried leaf weight (DW) per L in tea extract; <sup>^</sup>FLV = total flavonoids; n/a, not applicable

**Figure 2:**
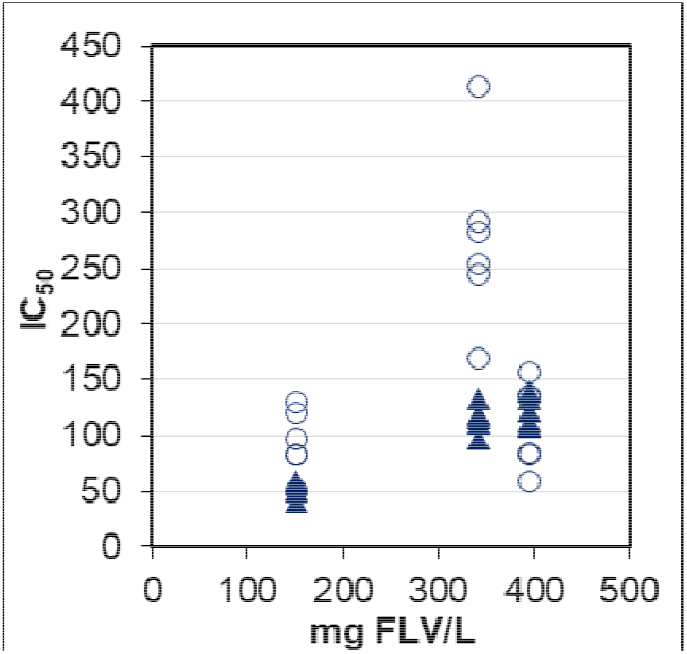
Spearman’s correlations between SAM, SEN, and MAL tea extract total flavonoids and IC_50_ for 2B6 (⍰) and 3A4 (▲); for 2B6, rho = 0.157 and *p* = 0.533; for 3A4, rho = 0.748 and *p* < 0.001.

### 4.2 Artemisia tea extracts differentially alter transcription of 2B6 and 3A4

After induction of HepaRG cells with ART, RIF, and the three Artemisia sp. extracts, qPCR showed that ART and RIF, as expected, significantly increased 2B6 and 3A4 transcription compared to the DMSO vehicle control (Fig. 3). Only A. annua SAM tea induced 2B6 expression compared to its water vehicle control; however, it was <20% that of ART despite having the same amount of ART (10 μM). A. annua SAM, on the other hand, significantly induced 3A4, about double that for ART at the same concentration (Fig. 3). Neither SEN nor MAL increased 3A4 or 2B6 transcription.

**Figure 3:**
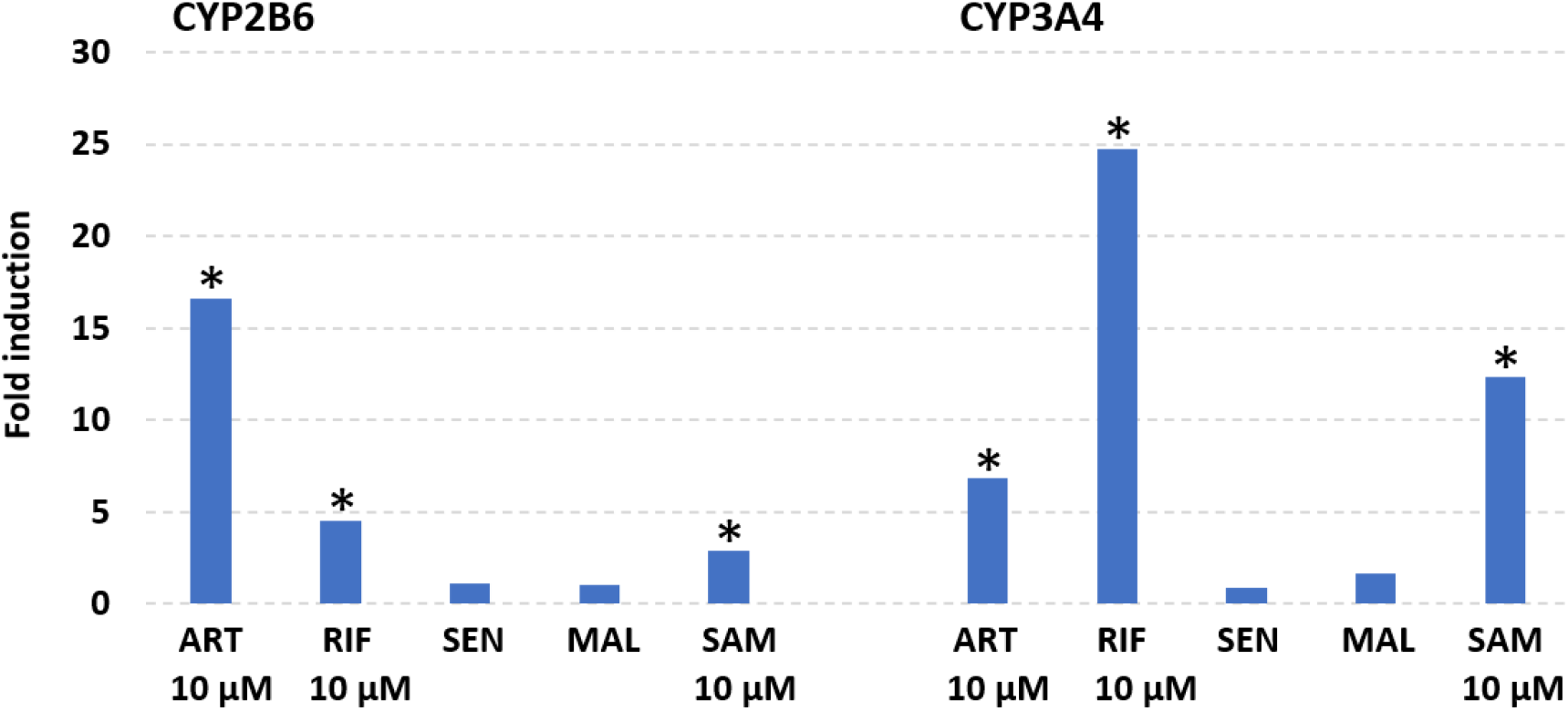
Fold change in 3A4 and 2B6 transcripts after 3-d induction of HepaRG cells with artemisinin (ART) and rifampicin (RIF) compared to DMSO solvent control; *A. annua* (SAM) and *A. afra* (SEN, MAL) and hot water extracts compared to water control. SAM and ART were at 10 μM ART; RIF was at 10 μM; SEN and MAL having little to no ART were tested at leaf dry mass equal to that of SAM at 10 μM ART; n = 3; Thermo Fisher cells. * = significant induction at fold induction > 2 relative to the vehicle (solvent) control, and at least 20% relative to the positive control; 2 biological replicate plates with n = 3 wells of cells/treatment/plate and 3 qPCR analyses/well.

When the A. annua SAM extract diluted to contain 10, 25, and 50 μM ART was added to HepaRG cells, CYP3A4 mRNA transcripts appeared to decrease from ∼2.0 to 0.2-fold as the A. annua SAM extract increased in concentration from 10 to 50 μM ART (Fig. 4), but the change was not significant between any of the dilutions (for 10 vs. 50 μM, ANOVA was *p* = 0.30). Although CYP2B6 mRNA transcripts increased with increasing ART in the SAM tea treatments, the change was not significant (ANOVA was p = 0.13 and 0.06 and for 10 vs. 25 and 50 μM, respectively). When compared to 10 μM ART, 2B6 and 3A4 transcripts were substantially less at all test tea concentrations containing 10-50 μM ART.

**Figure 4:**
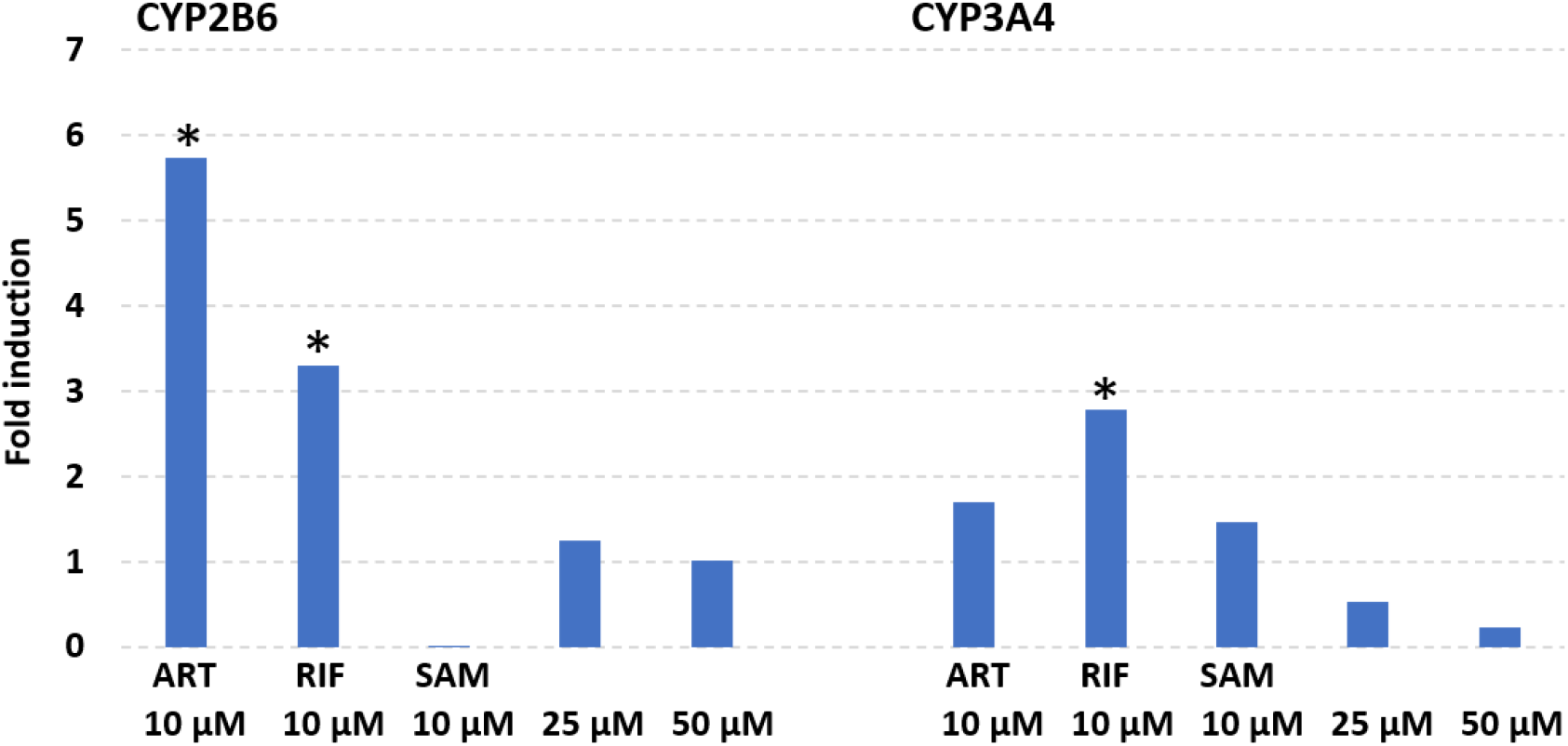
Fold induction of artemisinin (ART) and rifampicin (RIF) at 10 μM compared to DMSO solvent control; *A. annua* SAM hot water extracts at 10 μM, 25 μM, 50 μM ART compared to water control; n=3; Lonza cells. * = significant induction at fold induction > 2 relative to the vehicle (solvent) control, and at least 20% relative to the positive control; n = 3 wells of cells/treatment and 3 qPCR analyses/well.

**Figure 5:**
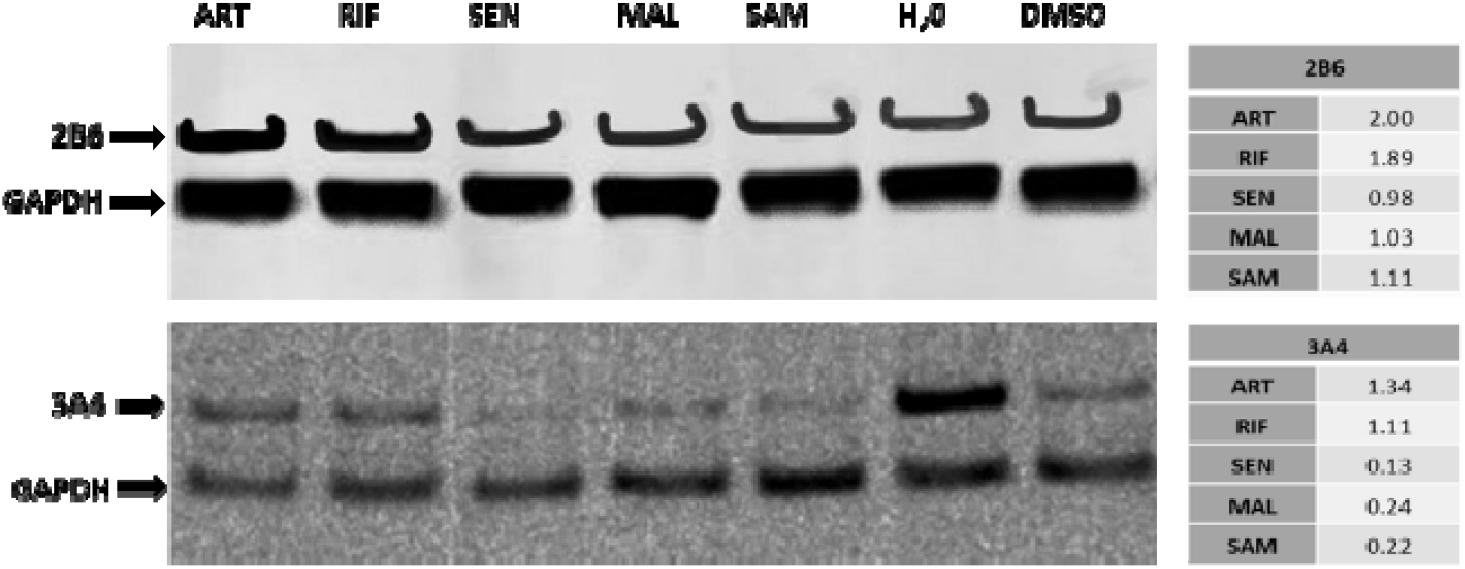
Representative Western blot analyses of 2-3 independent experiments showing 2B6 and 3A4 protein in HepaRG cells treated with artemisinin (ART) and rifampicin (RIF) compared to DMSO solvent control; and *A. annua* (SAM) and *A. afra* (SEN, MAL) hot water extracts compared to water control. SAM and ART were at 10 μM ART; RIF was at 10 μM; SEN and MAL having little to no ART were tested at a leaf dry mass equal to that of SAM at 10 μM ART; Thermo Fisher cells.

### 4.3 HepaRG cell responses differ per commercial source

Differences in RIF induction were noticed between separately induced batches of HepaRG cells (Fig. 3 and 4), which is consistent with some differential responses reported for these cells by the commercial vendors and even between different vials of cells, albeit each from the same lot # from the same vendor (Table 3; also see Table S1 in Supplemental Material and Discussion). Although Lonza and Thermo Fisher cells both responded to ART with an expected positive induction for both P450s, the Thermo Fisher cells (Fig. 3) provided a stronger response than Lonza cells, which for 3A4 expression was not significant (Fig. 4). Both cell sources provided a similar 2B6 response to RIF (Fig. 3 and 4), but for 3A4, Lonza cells (Fig. 4) had a lower response than Thermo Fisher cells (Fig. 3). Nevertheless, when all data from both cell sources were pooled together, both ART and *A. annua* SAM-treated cells induced 3A4 at 10 μM ART content (Table 3).

**Table 3.**
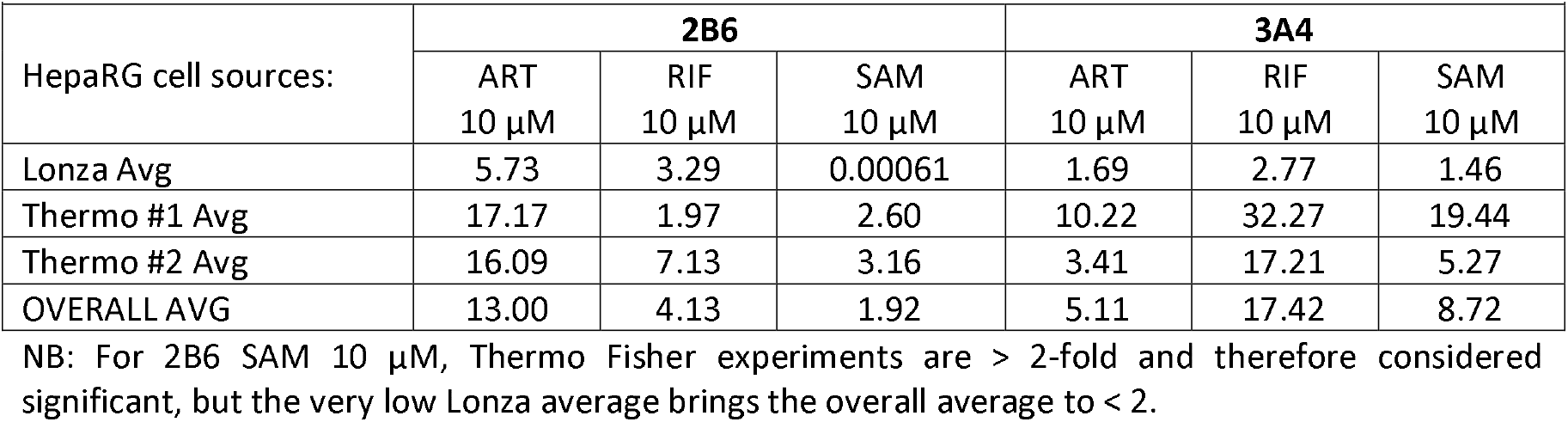
Comparative average in fold change between Lonza and Thermo Fisher HepaRG cells for ART, RIF, and SAM expression of CYP2B6 and CYP3A4.

|  |  |  |  |  |  |  |
| --- | --- | --- | --- | --- | --- | --- |
| HepaRG cell sources: | 2B6 |  |  | 3A4 |  |  |
| | ART<br>10 μM | RIF<br>10 μM | SAM<br>10 μM | ART<br>10 μM | RIF<br>10 μM | SAM<br>10 μM |
| Lonza Avg | 5.73 | 3.29 | 0.00061 | 1.69 | 2.77 | 1.46 |
| Thermo #1 Avg | 17.17 | 1.97 | 2.60 | 10.22 | 32.27 | 19.44 |
| Thermo #2 Avg | 16.09 | 7.13 | 3.16 | 3.41 | 17.21 | 5.27 |
| OVERALL AVG | 13.00 | 4.13 | 1.92 | 5.11 | 17.42 | 8.72 |

NB: For 2B6 SAM 10 μM, Thermo Fisher experiments are > 2-fold and therefore considered significant, but the very low Lonza average brings the overall average to < 2.

### 4.4 All Artemisia extracts reduced translation of 2B6 and 3A4 into protein

After induction of HepaRG cells with the *Artemisia* extracts, ART, RIF, and their respective solvent controls, Western blot analyses were run and analyzed for 2B6 and 3A4 protein levels (see Supplemental Material for all original Western blots). For 2B6, ART and RIF had about twice the protein of the DMSO solvent control, but all *Artemisia* extracts were essentially equal to the water control. ART and RIF induction showed slight increases in 3A4 compared to the DMSO solvent control. In contrast, *Artemisia* hot water extracts had about a fifth of the amount of protein of the water control for 3A4, suggesting post transcriptional inhibition. Thus, for both 2B6 and 3A4 there was no increase in protein after 3 days of induction with *Artemisia* extracts. Interestingly, there was significantly more 3A4 protein in the water control vs. the DMSO control; however, one can only compare *Artemisia* hot water extracts with their corresponding water control, and ART and RIF can only be compared with their corresponding DMSO solvent. Although for 3A4 it appears that more protein is produced with water vs. DMSO as solvent, this emphasizes the importance of choosing and normalizing to appropriate solvent controls. One cannot directly compare ART or RIF-induced protein levels to those of the *Artemisia* hot water extracts.

## 5. Discussion

Liver cytochrome P450s are one of the body’s first defenses against ingestion of toxic compounds and as such they metabolize most chemicals entering the body *per os*, with CYP3A4 being the most important drug-metabolizing enzyme (Guengerich, 1999). In comparison, CYP2B6, the major ART metabolizing hepatic P450, had been considered relatively minor with low expression (Mimura et al., 1993), but Stresser and Kupfer showed that there was considerably more CYP2B6 in the liver than previously thought (Stresser and Kupfer, 1999). CYP2B6 and CYP3A4 are the main hepatic P450s that metabolize ART (Svensson and Ashton, 1999).

Prior studies had shown that *A. annua* extracts inhibited activity of CYP3A4 and CYP2B6 in HLMs (Ashour et al., 2017; Desrosiers et al., 2020) and that even *A. afra* (SEN), with minimal ART content, also inhibited activity of both P450s (Desrosiers et al., 2020). This study confirmed those results in HLMs and included another cultivar, MAL, of *A. afra*. Although others showed that ART and dihydroART induced CYP2B6 and CYP3A4 transcripts (Xing et al., 2012), to our knowledge prior to this study the role of the traditional tea infusions on gene expression and translation of these two P450s remained unknown.

The present study used traditional tea infusions and showed that in contrast to ART alone, the infusions of *A. annua* and *A. afra* not only inhibited enzymatic activity of CYP3A4 and CYP2B6, but also inhibited ART-driven induction of CYP2B6 mRNA transcripts. Zhang et al. (Zhang et al., 2020) previously showed about a five-fold induction of CYP3A4 compared to RIF when ART increased from 1 to 100 μM. However, we saw no significant change with increasing ART-delivered *A. annua* SAM. Furthermore, post transcriptional analysis by Western blot revealed no change in 2B6 and ∼80% decrease of CYP3A4 protein levels by all *Artemisia* hot water extracts, even SAM containing a high level of ART, suggesting *Artemisia* extracts inhibited ART-driven expression of CYP3A4 protein at the post-transcriptional level since qPCR had shown an increase in CYP3A4 transcripts with 10 μM ART.

Another observation was the appearance of considerably more CYP3A4 protein expression in water than in DMSO. Water is a solvent more akin to the milieu for an orally digested drug than DMSO and thus better represents *in vivo* metabolism. While one cannot directly compare analyses done in DMSO with those done in water, the decrease in apparent CYP3A4 protein with DMSO has some precedence in the literature. For example, Hoekstra et al. (Hoekstra et al., 2011) showed that DMSO reduced total protein by ∼60% in HepaRG cells. There are other reports of such solvent variations on CYP3A4 protein levels; Zhang et al. showed that the solvent, CITCO, used in studies involving constitutive androstane receptor (CAR) increased CYP3A4 protein levels by ∼30% compared to DMSO (Zhang et al., 2020).

We also observed that increasing the amount of *A. annua* SAM tea infusion from 10 μM to 50 μM ART increasingly inhibited transcription of CYP3A4, but there was a reverse effect on CYP2B6. Although neither result was statistically significant, results may suggest a combination effect deserving of more study. *A. annua* SAM contains about 16 mg of ART/g of dried leaves, so it is likely that the lower concentration of ART (10 μM) in the tea infusion was relatively high compared to other phytochemicals that could have inhibited CYP3A4 transcription. Although the ratio of ART to other phytochemicals remained constant, at the higher (50 μM) concentration, the non-ART constituents may have reached an antagonistic concentration in the less dilute tea infusion that subsequently interfered with CYP3A4 transcription. There are many phytochemicals present in these *Artemisia sp*. that reportedly inhibit CYP3A4 activity and could possibly account for the response of CYP3A4 in this study including: AB (Cai et al., 2017; Desrosiers et al., 2020); α-thujone, 1,8-cineole, borneol, limonene (Zehetner et al., 2019); chrysoplenetin (Wei et al., 2015); quercetin, rutin, and scopoletin (Ashour et al., 2017; Desrosiers et al., 2020). Many occur in both *A. annua* and *A. afra* (Brown, 2010; Liu et al., 2009; Sun et al., 2015). However, relative phytochemical concentration effects on CYP3A4 as well as CYP2B6 transcription remained to be studied.

Although differences in RIF induction were noticed between separately induced batches of HepaRG cells, they are consistent with the use of cells sourced from either Thermo Fisher or Lonza. Lonza reports HepaRG RIF induction of CYP3A4 at ∼9-fold (https://bioscience.lonza.com/medias/HepaRG-CYP-Induction-chart?context=bWFzdGVyfHBpY3R1cmVQYXJrTWVkaWF8MTI5NzM4fGltYWdlL2pwZWd8cGljdHVyZVBhcmtNZWRpYS9oZWMvaDEyLzkwNzk0OTEwNjc5MzQuanBnfDVlMTkxNWFjMTA3YTE3OGNhOTBiMjc1ZmJhMDIxYzEyODI2MzdiZGIyZTVkNmQ5ZTRlNDMxMWNlNmUwOTY2YTk, accessed May 6, 2022) compared to our induction of ∼2.8-fold. Thermo Fisher HepaRG cells are reported to induce ∼20-fold by RIF (https://www.thermofisher.com/us/en/home/industrial/pharma-biopharma/drug-discovery-development/adme-tox/hepatic-systems/heparg-cells.html, accessed May 5, 2022), while our data show an average induction level of about 25-fold induction for CYP3A4. It is thus clear that the level of RIF induction of CYP3A4 varies substantially between HepaRG sources. Indeed, even from a single commercial cell source there are variations in level of RIF induction (Table S1). Nevertheless, internal solvent controls normalize data for each cell set and the other positive control, ART, also induced CYP3A4; and as the pooled averages from all our induction experiments showed, ART, RIF, and *A. annua* SAM all induced CYP3A4 at 10 μM.

Reported inhibitors of CYP2B6 activity include: borneol, limonene, linalool (Zehetner et al., 2019); quercetin, AB (Desrosiers et al., 2020); camphor, and 1,8-cineole (Seo et al., 2008). Absence of reported inhibition by individual phytochemicals does not necessarily indicate lack of inhibition by an extract against both P450s because studies do not analyze all of the phytochemicals found within a complex extract. There also could be both antagonistic as well as synergistic effects. Conflicting results for some phytochemicals are reported. For example, in contrast to Desrosiers et al. (Desrosiers et al., 2020), Cai et al. (Cai et al., 2017) reported there was no inhibition of CYP2B6 by AB. However, the former used P450-Glo with HLMs while the latter used HLMs with CYP2B6- and CYP3A4-specific substrates measured by LCMS. In another instance, Zhang et al. (Zhang et al., 2020) did not measure enzyme activity and instead used CYP2B6 and CYP3A4 luciferase reporter gene plasmids transfected into HepG2 cells and a Promega kit that converts luminometer readings to gene expression levels; they also suggested that the Cai et al. (Cai et al., 2017) discrepancy could be attributed to differences in study design and determination of activity. Such discrepancies in data comparisons are challenging and remain to be resolved using uniform comparative methods. Further, when using herbs, people do not take isolated compounds. They consume a complex extract or infusion or the whole herb (Na et al., 2011). Indeed, it is anticipated that although there may be individual compounds that are pharmacologically active, there are likely other compounds present that modulate enzymes that provide specific metabolic responses, e.g., liver P450s.

## 6. Conclusions

This is the first study that compared ART to *Artemisia sp*. hot water extracts for their effects not only on CYP2B6 and CYP3A4 enzymatic activity, but also their transcription and translation into enzymatically active protein. These results suggest that continued *per os* consumption of traditionally prepared *Artemisia* tea infusions likely may provide a consistent daily dose of ART with no anticipated losses from CYP2B6 and CYP3A4 metabolism. This is important for establishing longer-term understanding of ART serum concentrations after multiday *per os* consumption of *Artemisia* tea infusions. This study also offers caution for *Artemisia* use when combined with other drugs because inhibition, especially of CYP3A4, could lead to altered serum drug concentrations with possible adverse physiological effects. Future work will focus on bio-guided fractionation and identification of specific phytochemicals in these plants that are responsible for the observed effects on CYP2B6 and CYP3A4 and interactions of *Artemisia* with various commonly prescribed drugs.

7.

## Glossary

AB: arteannuin B
ACT: artemisinin combination therapy
ART: artemisinin
CYP: cytochrome P450
DLA: dried leaf *Artemisia*
FLV: flavonoids
HLM: human liver microsome
MAL: *Artemisia afra* Malawi cultivar
PHH: primary human hepatocytes
RIF: rifampicin
SAM: *Artemisia annua* US cultivar
SEN: *Artemisia afra* Senegal cultivar

## 8. Acknowledgements

The authors thank Worcester Polytechnic Institute and La Maison d’Artemisia for jointly funding a Global Fellowship for NF Kane along with some project supplies. Prof. BH Kiani was partially supported by an American Association of University Women Fellowship. Thanks to Profs. J. Arguello and R. Page at WPI for use of equipment for PCR and Western analyses. Award Number NIH-2R15AT008277-02 to PJW from the National Center for Complementary and Integrative Health funded this study. The content is solely the responsibility of the authors and does not necessarily represent the official views of the National Center for Complementary and Integrative Health or the National Institutes of Health.

## 9. Authorship Contributions

NF Kane – planned experiments, conducted experiments, wrote the manuscript, edited manuscript

MR Desrosiers – planned experiments, conducted experiments, edited manuscript

BH Kiani – conducted experiments, wrote manuscript, edited manuscript

MJ Towler – conducted experiments, edited manuscript

PJ Weathers - obtained funding, planned experiments, supervised research, wrote manuscript, edited manuscript

## SUPPLEMENTAL MATERIAL

### Modified total flavonoid assay

For methylene chloride extracts of dried leaves, total flavonoids were determined by adding 1 mL of 1% AlCl3 in methanol to test tubes containing dried aliquots of extract (typically 2.5 to 5 mg DW), incubating room temperature for 30 min, and reading the absorbance at 415 nm. Standards of quercetin were treated similarly. For hot water extracts, aliquots of tea were diluted with methanol to a volume of 500 μL, and then 500 μL 2% AlCl_3_ in methanol was added. Standards of quercetin had a volume of water equivalent to the aliquot of tea added, plus methanol to a volume of 500 μL, and then 500 μL 2% AlCl_3_ in methanol. Samples were incubated and analyzed as described previously.

### HepaRG cell proliferation

The cells were cultured for 3-d before starting the 3-d induction assay. Photos were taken at different times to check cell morphology and integrity. At 24 h after plating, cells were already attached and appeared in small-differentiated colonies. At 72-96 h cells formed a monolayer and were organized in clusters with well-delineated trabeculae with many bright canaliculi-like structures (Figure S1).

**Figure S1.**
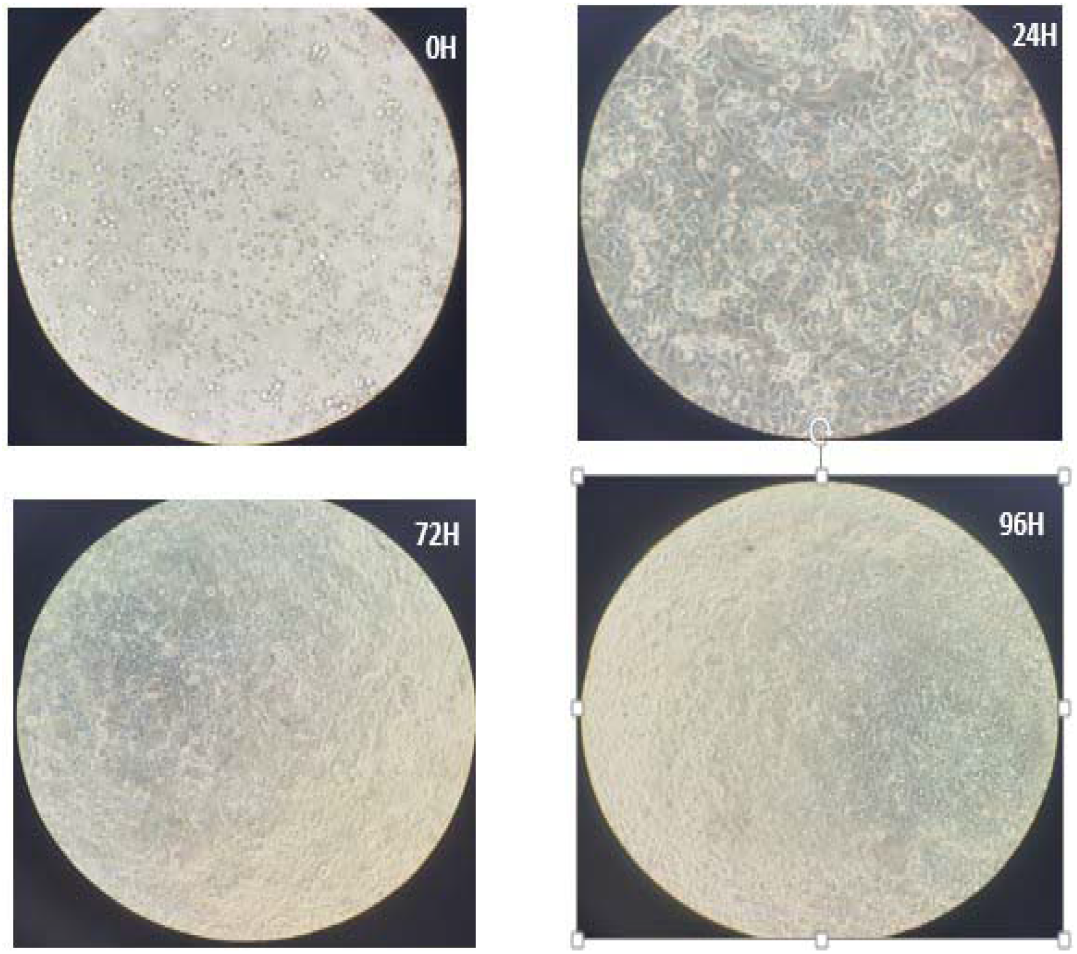
HepaRG cells during the 72-h induction process.

### Original Western blot images

**Figure S2:**
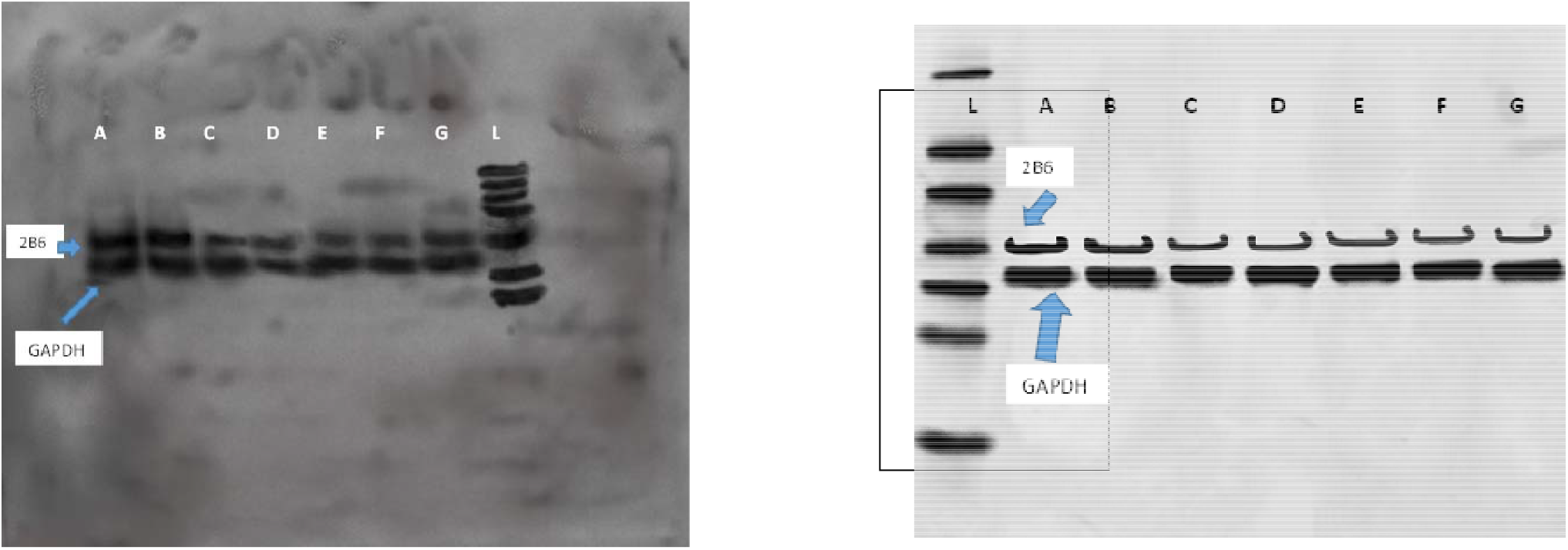
2B6 blots each from a separate HepaRG experiment.

**Figure S3.**
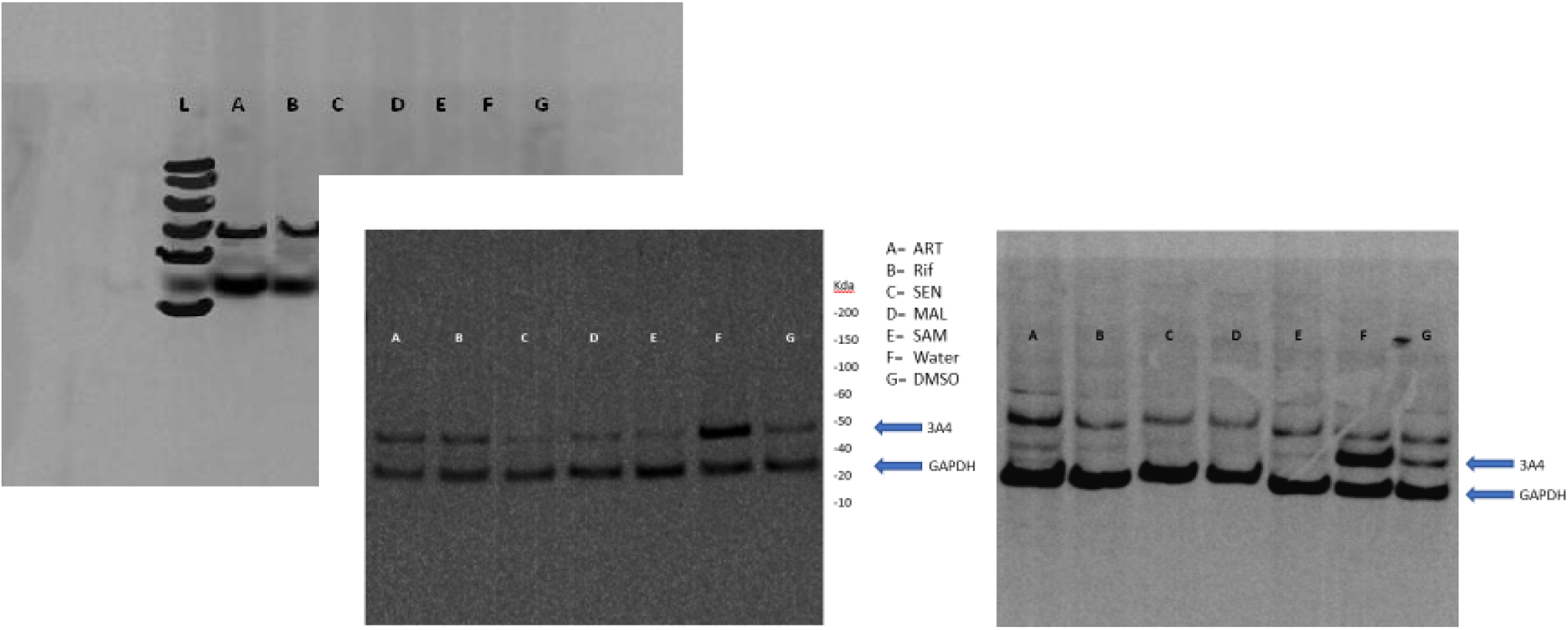
3A4 blots each from a separate HepaRG experiment.

### Differential responses to RIF from different vials of HepaRG cells from a single vendor (Lonza)

Two preliminary HepaRG induction experiments were conducted to determine if there was any evidence of induction and its relative level using P450-Glo assays. HepaRG cells from Lonza were used for the P450-Glo assays of 2B6 and 3A4 activity indicative of P450 activity post-induction. Collagen I coated 96-well plates were used, and cells were plated at a seeding density of ≅100,000 cells/well. Cells were cultured for 3 d, followed with a 3-d induction using a medium containing RIF (10 μM) as positive inducer and compared to the solvent control (0.1% DMSO). After the induction, cells were washed twice with PBS with 3 mM salicylamide. To 2B6 wells, 50 μL of luciferin-2B6 was added and cells were incubated at 37°C for 120 min. To 3A4 wells, 50 μL of luciferin-IPA was added and cells were incubated at 37°C for 60 min. To each well, 50 μL of detection reagent was added and mixed briefly using a plate shaker. The plate was equilibrated at room temperature for 20 min, after which luminescence was measured using a plate reader. **Table S1** shows response of 2B6 and 3A4, respectively, in response to RIF at 10 μM.

**Table S1.** Differential response of 2B6 and 3A4 to different vials of Lonza HepaRG cells after induction with 10 μM rifampicin.

| **Table S1.** Differential response of 2B6 and 3A4 to different vials of Lonza HepaRG cells after induction with 10 μM rifampicin. | | |
| --- | --- | --- |
| **Experiment #** | **Fold change in P450 post P450-Glo assay** |  |
|  | **2B6** | **3A4** |
| 1 – RIF (MRD) | 2.5 | 11.5 |
| 2 – RIF (NFK) | 5.0 | 5.3 |

